# Volumetric fluorescence imaging in a human eye model by oblique scanning laser ophthalmoscope (oSLO): a feasibility study

**DOI:** 10.1101/804237

**Authors:** Wenjun Shao, Weiye Song, Ji Yi

## Abstract

Fluorescence retinal imaging, such as fluorescein angiography, indocyanine green angiography, and autofluorescence imaging, are valuable tools in ophthalmology and vision science. However, these clinical imaging modalities provide *en face* view of the retina, with limited capability to discriminate retinal layers over a large field-of-view (FOV). We recently developed a novel retinal imaging method, oblique scanning laser ophthalmoscopy (oSLO), to provide volumetric retinal fluorescence imaging without any depth sectioning. OSLO breaks the coaxial alignment of the excitation and detection, to produce a cross-sectional view on retina using the natural ocular optics. In this paper, we demonstrated oSLO in a realistic human eye model and showed the feasibility for future *in vivo* human retinal imaging. A new optical design was implemented to significantly simplify our previous oSLO systems. We overcame the limitation by the small numerical aperture (NA) of the human eye, by integrating a pair of cylindrical lens in the remote focusing system. We experimentally showed that the current setup can achieve a FOV of ∼3×6×0.8 mm^3^, and the transverse and axial resolutions of 7 and 41 µm, respectively. The capability of volumetric fluorescence imaging over a large FOV in the human retina could lead to new clinical imaging paradigms for retinal diseases.

## 1. Introduction

The development of modern optical retinal imaging methods has revolutionized the clinical practice of ophthalmology and vision science. Each technique has its unique strength and is best suited for certain retinal pathologies [1–5]. Among them, optical coherence tomography (OCT) provides 3D structural images of the retina and choroid with micrometer-scale resolution. OCT angiography (OCTA) has emerged as an important tool in imaging 3D microvasculature over a large field-of-view (FOV) non-invasively [6]. One major limitation of OCT is that it detects the scattering signal, and thus is insensitive to fluorescence contrasts [7]. On the other hand, fluorescence-based imaging methods, such as fluorescein angiography (FA), indocyanine green angiography (ICGA) and autofluorescence imaging are valuable in providing other functional information. Conversely, all the fluorescence-based imaging methods in clinics have limited capability in volumetric retinal imaging.

One common approach for all the fluorescence-based imaging methods is scanning laser ophthalmoscope (SLO) [8]. By using a flying laser beam, SLO permits imaging at low light intensities and avoids the amount of unwanted light exposure outside the focal volume. To improve the resolution and enhance the contrast, confocal gating was used in SLO to block the diffusive light [9]. In order to observe finer retinal structures, parallel confocal line was adopted in SLO systems, and lateral resolution of 3.91 µm for human eye was reported [10]. Although the lateral resolution of SLO-based approaches has gained considerable improvement, the depth sectioning ability is restricted by the depth of focus of the excitation light. Even with an optimized pinhole size, the axial resolution for the human eye is 200-250 µm due to the aberration when the pupil is fully dilated [11,12]. Apparently, this resolution is insufficient to provide depth discrimination within retina for volumetric imaging [13].

With the application of adaptive optics technique, the way we image the retina has been changing [14]. As adaptive optics corrects the wavefront error caused by the aberration of the imperfect optics in the human eyes, both the lateral and axial resolution of an adaptive optics scanning laser ophthalmoscope (AOSLO) could approach the diffraction limits. The lateral resolution of AOSLO is revealed to be 2 - 5 µm, and the axial resolution was 37 – 84.2 µm for the human retina [15–23]. Even though the axial resolution is improved, the volumetric imaging by AOSLO in human retina is plagued by the need for depth-sectioning and small FOV, typically within 1° - 2°.

To enable volumetric retinal imaging for fluorescence contrasts, we recently developed a novel method, called oblique scanning laser ophthalmoscope (oSLO). The novelty of oSLO is that it breaks the coaxial alignment of the emission and excitation in the conventional SLO, by using the asymmetric, off-axis excitation and detection. As the off-axis excitation light creates an angle with the optical axis of the human eye, a tilted 2D cross section of the human retina can be illuminated and imaged. By sweeping the excitation light, the volumetric retinal imaging can be achieved without depth sectioning. We have previously demonstrated the success of oSLO in rats and mice *in vivo* over ∼30° FOV with sufficient depth resolution to resolve the stratified microvascular plexus [24,25]. As the excitation and emission light travel in two separated light paths in the previous design, two synchronized galvanometer mirrors were used for scanning and descaning, which makes the optical system bulky and complex in alignment. To overcome this disadvantage and make the oSLO suitable for the practical application in the clinic, a new optical design was implemented, resulting a compact system setup. With the novel oSLO system, we demonstrated the feasibility of oSLO for human retinal imaging using a realistic eye model. We overcame the challenge of small NA ∼0.2 for human ocular optics and characterized the later and depth resolution of 7 µm and 41 µm, respectively, and a ∼10° FOV.

## 2. Experiment setup and methods

### 2.1 The overall design of oblique scanning laser ophthalmoscopy for the human eye

The schematic of oSLO for the human eye is shown in Fig. 1(a). The 3D modeling in Fig. 1(b) was used to validate the feasibility of the overall optical and mechanical design. Figure 1(c) is a photograph of the actual oSLO system. The setup is compact that can easily be transformed for clinic used in the future.

**Fig. 1.**
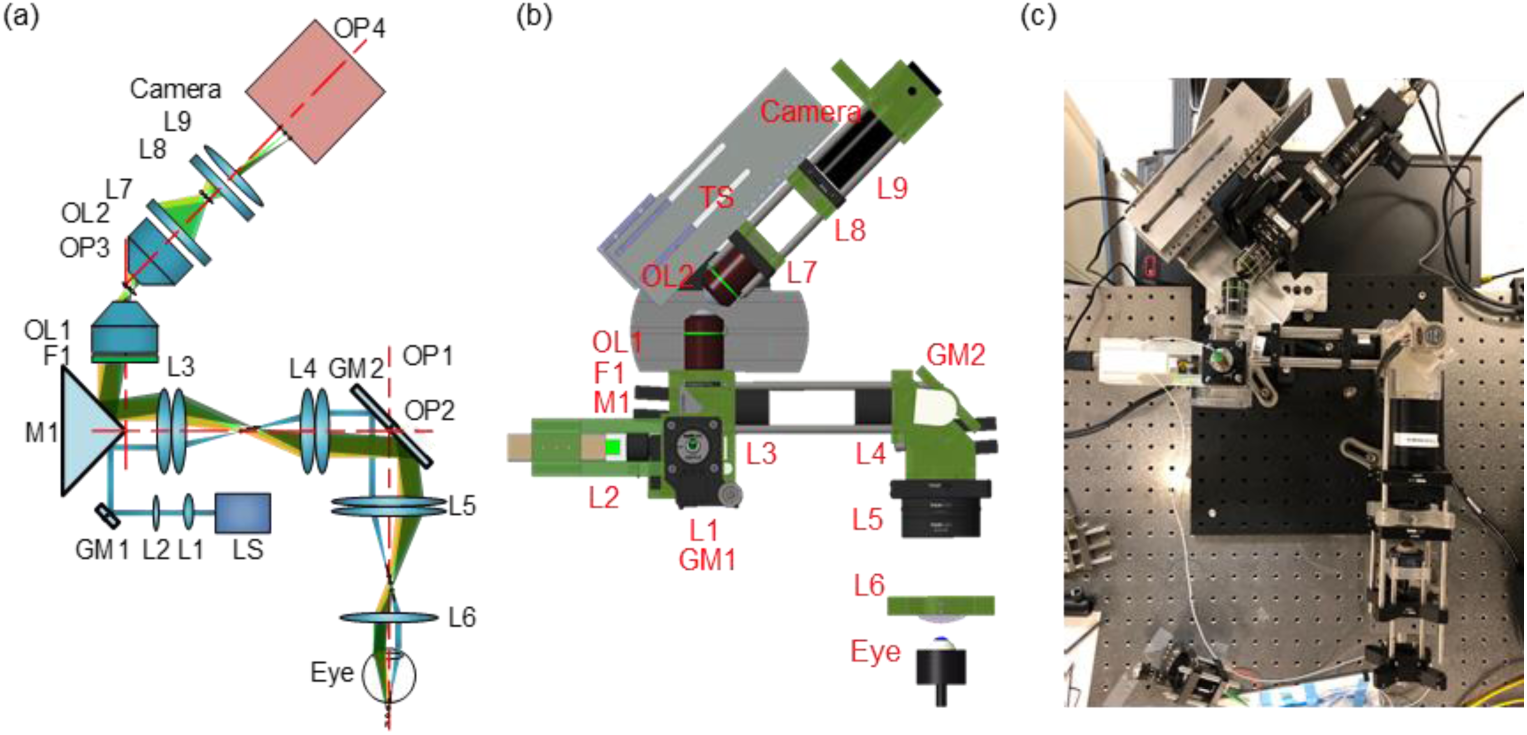
Experiment setup of oSLO for the human eye. (a) The system schematic. L: lens; OL: objective lens; F: filter; M: mirror; GM: galvanometer mirror; LS: light source, OP: optical axis, TS: translation stage. (b) The 3D modeling of oSLO system. (c) The photography of the actual oSLO system.

The light source (LS) is a 488 nm laser with 30 mW maximum power. The laser was first coupled into a single-mode fiber and collimated by an *f* = 10 mm lens (L1). To maintain a confocal alignment of the excitation and detection path over a large FOV, we used a focus tunable lens (L2) to dynamically adjust the focus position of excitation light during imaging. A galvanometer mirror (GM1) was used to steer the excitation light to form a raster laser line. A right-angle prism mirror (M1) was used to separate the excitation from emission light, which was marked in different colors (blue for excitation light; green, yellow and green for emission light). On one side of the mirror, the excitation light was first relayed to the slow scanning galvanometer mirror (GM2) by 1:1 relay lens (L3: *f* = 50mm, L4: *f* = 50 mm), and further to the pupil entrance by 2:1 relay lens (L5 *f* = 50 mm and L6: V40LC, Volk lens, *f* = 25 mm). To form an oblique light sheet in the retina, the excitation light was offset from the optical axis (OP2). The offset distance is ∼7mm, resulting a ∼3.5mm offset at the pupil entrance. In this case, the excitation light entered the human eye from the edge of the pupil, resulting an oblique angle of the excitation light sheet to be ∼10° [24]. GM2 was used to scan the excitation light so that the tilted light sheet could sweep through the retina, and also descan the emission light. At the same time, the fluorescence emission from the retina was mapped back through the relay lenses (L3-L6), and turned to the back pupil plane of OL1 (UplanFL N 20 × /0.75) by the prism mirror (M1). An emission filter F1 was placed in front of objective lens OL1 for wavelength within 500-550 nm. A stationary and tilted image could then be formed after OL1. As the image after the objective lens OL1 was tilted, the optical axis OP4 of the final imaging system (OL2: UplanFL 20 × /0.5, L7: *f* = 50 mm, L8: *f* = 10 mm and L9: *f* = 50mm) was also tilted to remotely focus the tilted image onto the camera sensor (BFS-U3-51S5M-C, Point Grey). The intersection angle of OP3 and OP4 could be precisely adjusted by a mechanical stage (TS) with 3 degrees of freedom, as can be seen in Fig. 1(c). The choice of this angle was to keep a balance between FOV and emission light collection efficiency.

As shown in Fig. 1(c), the human eye model was placed at the front of L6 with a distance of ∼22mm, allowing a pleasant working distance for the human eye. The power of the laser beam projected on the cornea was limited within 0.5 mW, which was below the maximum permissible exposure (MPE) of lasers established by the American National Standards Institute and other international standards [26,27]. The exposure time of the camera was set to 10 ms, permitting a frame rate of 100 FPS.

### 2.2 Optics design for low NA oSLO by using cylindrical lens

Figure 2(a) is a simplified sketch from Fig. 1(a). Object 1 represents the retina. The conjugate image of the retina (image 2) is between OL1 and OL2. The image 3 is on the camera sensor. Due to the small NA of the human eye, the excitation light sheet has a small angle of α ∼10° with respect to the optical axis. The conjugate image2 has an angle β with respect to the optical axis. The transverse magnification from 1 to 2 was designed to be *M_X_* = 0.2, to increase the angle β to be ∼41° by the Scheimpflug condition [28]:

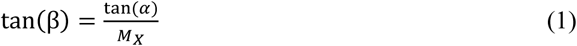

**Fig. 2.**
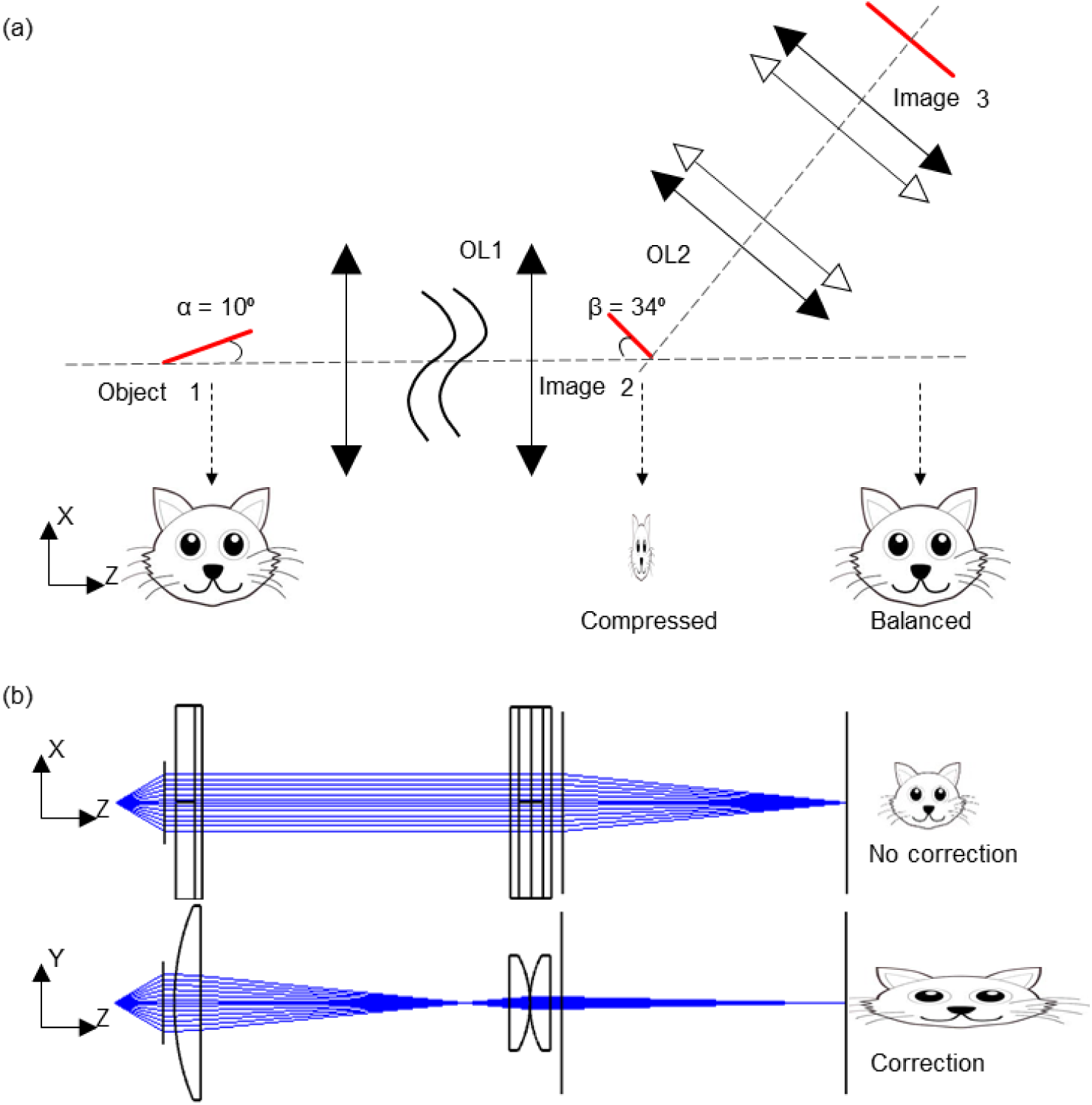
The schematic of the low NA oSLO by using cylindrical lenses. (a) The optical design for the correction of both the angle and aspect ratio of the image. (b) The correction of the compressed image by the unidirectional optical power of the cylindrical lens.

The corresponding axial magnification is then 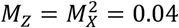[29]. As a result, image 2 was heavily compressed in the axial direction and asymmetric in its two dimensions. A kitten face is used in Fig. 2(a) to vividly illustrate the unbalanced magnification in two dimensions.

To achieve a unidirectional magnification, we applied an anamorphic telescope composed of two groups of cylindrical lenses (marked with hollow arrows) to magnify the depth dimension by 5x and have no power in lateral dimension, as shown Fig. 2(a). As a result, the overall magnifications from object 1 to image 3 were corrected to be consistent in its two dimensions, about 1:1.

### 2.3 OCT imaging

A human eye OCT system described from our previous publications [30,31] was used to characterize the physical dimension of the retina phantom and provide OCT angiography (OCTA) as a simulation image for oSLO in human FA. The system used 800-880 nm laser as light source. The exposure time for each A-line is 19.1 µs and 9.1 µs for OCT and OCTA, respectively. To acquire an OCT image, triangular waves with 50% duty cycle and a ramping voltage were used to control the fast and slow galvanometer, respectively. Each B-scan contains 512 A-lines in the forward or backward scanning direction. One OCT volume contains 256 B-scans. For OCTA, the control signal duty cycle of the fast galvanometer for OCTA is 80% and each B-scan contains 320 A-lines only in the forward direction. Three repeated B-scans were acquired at the same location in the y-direction. One OCTA volume contains 320 B-scans. To generate OCT and OCTA image, the raw data were normalized by the light source spectrum first, then several preprocessing steps were performed including removing the DC spectral component, λ-k resampling and digital dispersion compensation [32]. Then a Fourier transform on the interferogram generated OCT images. For OCTA, we took a similar strategy from the split spectrum strategy as in SSADA [33], but used complex signals for angiography contrast [34]. The FWHM of the Gaussion window in our split spectrum processing was 0.10 µm^-1^ in *k* spectral space, resulting an axial resolution of ∼26 µm. The axial resolution of structural OCT was ∼5.6 µm. The lateral resolution for OCT and OCTA was defined by the incident beam width which was estimated to be ∼8.2 µm and ∼10.2 µm, respectively.

### 2.4 The design of the human eye model

We used a realistic eye model to test the feasibility of oSLO on human retina. The schematic is shown in Fig. 3(a)-3(b), and it was modified from a commercial product (OEM-7, Ocular Instruments, Bellevue, WA). The model consists of the cornea, crystalline lens, aqueous humor, vitreous humor, and artificial retina. As depicted in Fig. 3(a), the retina was emulated by a spherical shell of 0.5% agarose gel attached to the bottom of the eye model. Fluorescent microspheres with 3.1 µm diameter were embedded in the gel to characterize the imaging resolution. To prevent the agarose gel from detachment, a 3D printed plastic fixture was used to hold the gel in place. On the bottom of the plastic fixture, a 6 mm diameter clearance hole was designed for imaging, as shown in Fig. 3(b).

**Fig. 3.**
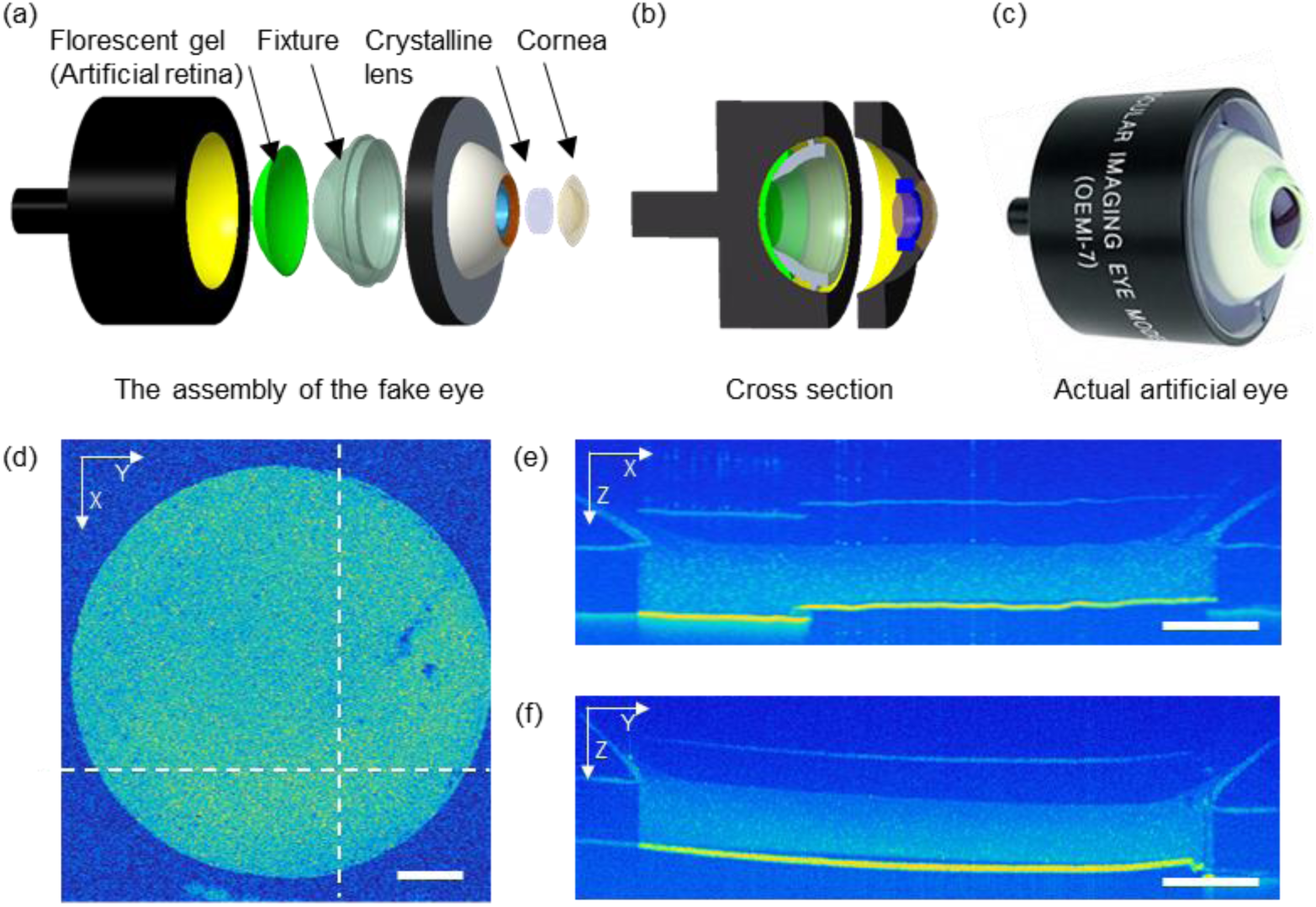
The design of the human eye model and preparation of the agarose gel as an artificial retina. (a) The 3D assembly and the inner structure of the human eye model. (b) A cross section of the 3D model of the human eye. (c) Photography of the actual human eye model. (d-f) The X-Y, X-Z, and Y-Z cross-sectional images captured by an OCT device. Bar = 1 mm.

To measure the physical dimension of the artificial retina, OCT imaging was performed on the eye model. The theoretical resolutions in axial and transverse direction for our OCT setup were ∼5.6 µm and ∼10.2 µm [30], respectively, providing sufficient measurement accuracy. Figure 3(d)-3(f) were two OCT B-scans, and the *en face* projection of the artificial retina. The boundary of the clearance hole and the entire thickness of the retina can be visualized. The diameter of the clearance hole is used to calibrate the transverse dimension. The thickness of the artificial retina was measured to be ∼0.8mm.

### 2.5 Data acquisition and image processing

The synchronization of the camera trigger, galvanometer GM1, and GM2 was modified from our previous publications [24]. Briefly, the camera was triggered to acquire a 2-D B-scan fluorescent image every 10 ms. In every trigger period, the fast scan (GM1) was controlled by triangular waves with 2 ms period and 50% duty cycle to create a scanned light sheet. At the end of every trigger period, the slow scan (GM2) was driven by a square wave to move an incremental step forward. When the slow scan was sweeping across the retina, the focus for the excitation was adjusted by the tunable lens with a ramping wave, allowing confocal alignment of excitation and detection. The slope of the ramp wave was determined by experimentally measuring the focal shift along the whole trip of the slow scan. In order to recover the physical geometry, the 3-D matrix was sheared by 10° using affine transformation to correct the oblique illumination angle.

## 3. Results

To verify whether this optical design could unfold the compressed depth information and balance the magnifications in the transverse and axial direction, an imaging target was fabricated, as shown in Fig. 4(a). The fluorescent microspheres of 3.1 µm diameter were immobilized in 0.5% agarose gel, which was sandwiched between the four glass slides forming three layers of agarose gel. The thickness of every layer is 1.2 mm. To image this imaging target, the human eye model in Fig. 1(a) was replaced by a lens with *f* = 25mm. Figure 4(a) and (b) are two cross-sectional images captured without and with cylindrical lens (L7 and L8), respectively. There were three agarose gel layers from left to right side of the image. The image of the microspheres is sharp and clear at the left-most layer while it gradually becomes blurred toward the right-most layer. This is because the left-most layer of the agarose is within the depth of focus of the excitation light while the other two layers are out of the depth of focus. The depth information in Fig. 4(a) was dramatically compressed. On the contrary, the depth dimension in Fig. 4(b) was magnified by using cylindrical lenses. As can be seen from the comparison in Fig. 4(a) and (b), the width of the phantom image in Fig. 4(b) is approximately 5 times that of the image in Fig. 4(a), which validated our optical design. The overall magnifications in the lateral and axial direction are consistent to be ∼0.9. As the lens in front of the phantom has a different focal length with the realistic eye, the magnification is slightly different from the previous calculation. But it still serves as proof of the aspect ratio correction in our optical design.

**Fig. 4.**
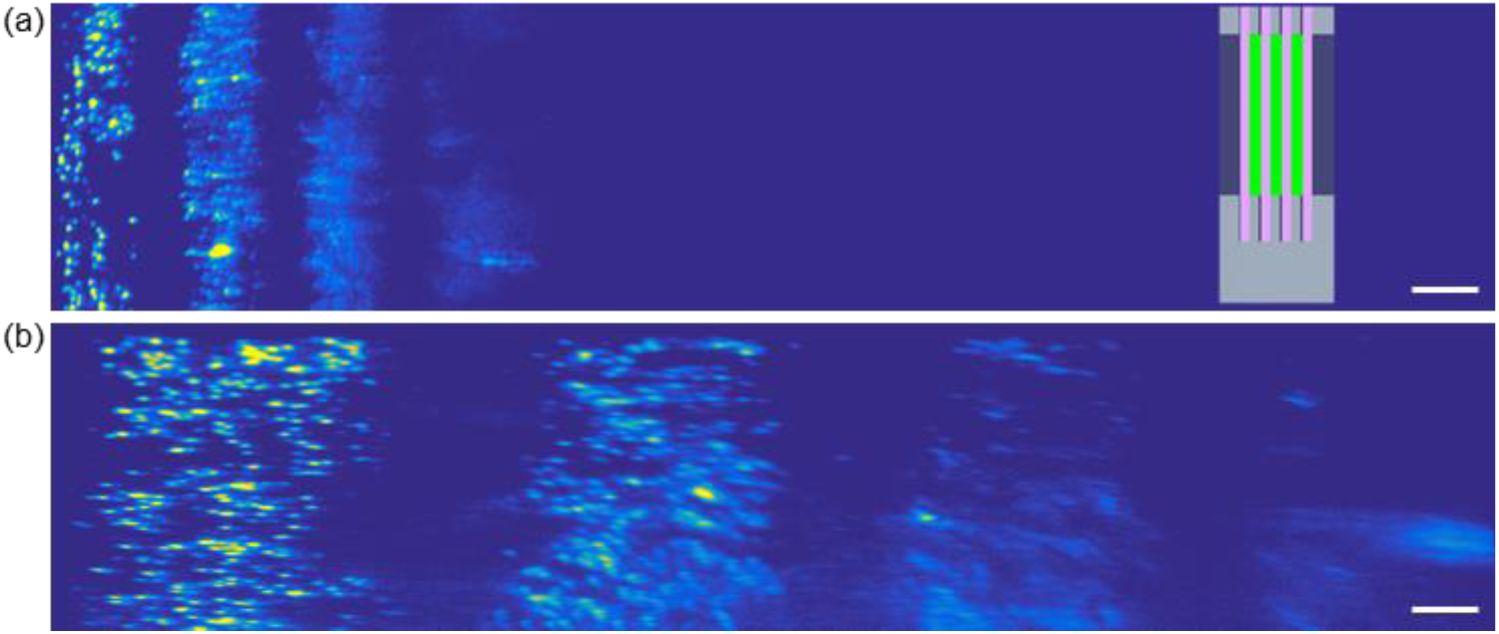
The verification experiment for the optical design based on cylindrical lens. (a) 3D model of the image target with three fluorescent layers. (b) A cross section of 3D model showing the inner structure of the image target. (c) Cross-sectional image captured without cylindrical lenses. Bar = 0.25 mm. (d) Cross-sectional image captured with cylindrical lenses. Bar = 0.25 mm.

In order to experimentally characterize the resolution and the feasibility of the proposed oSLO, the artificial retina adhered in the bottom of the human eye model was imaged.

Figure 5 shows the representative oSLO images acquired from the artificial retina. The maximum intensity projections on three planes were displayed in Fig. 5(a)-5(c). The FOV covered half of the clearance hole of the fixture, achieving ∼6 × 3 mm^2^ area. The shape of the curve in Fig. 5(a) was consistent with that of the OCT result shown in Fig. 3(f). The length of the FOV is sufficient to cover the macular area, typically ∼5-6 mm in diameter. The FOV on the fast axis is currently limited by the size of the camera sensor size. Figure 5(d)-5(f) are the zoomed-in views from Fig. 5(a)-5(c). The axial resolution appeared consistent throughout the depth of 0.6-0.7 mm, which is sufficient to penetrate the entire retina for a typical thickness <0.5 mm. Meanwhile, the lateral resolutions were well maintained over the entire FOV.

**Fig. 5.**
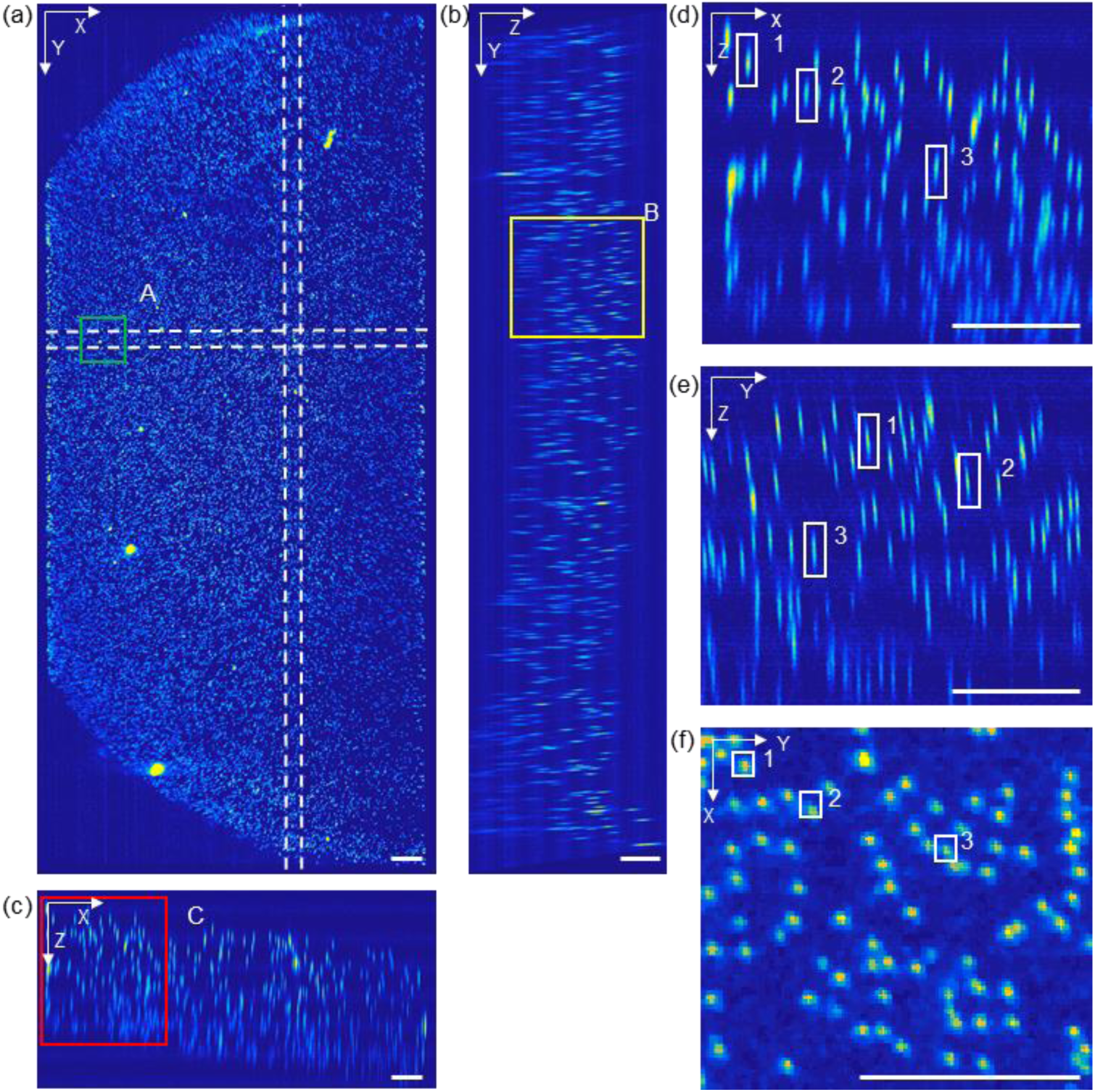
Feasibility and resolution characterization of oSLO for the human eye. (a-c) Maximum intensity projection of the 3-D volumetric fluorescent image in X-Z, Y-Z and X-Y planes. (d-f) Zoomed-in view of the square area A, B and C in panel (a-c), respectively. Three cross-sectional fly-through videos are included in the supplemental material. Bar = 0.25 mm.

Due to the small diameter, the fluorescent beads served as point sources to characterize the point spread function (PSF) of our oSLO system. Three representative beads labeled in Fig. 5(d)-4(f) were randomly selected to characterize the 3D PSF. Maximum intensity projection was applied to each of the three beads along x, y and z directions with the corresponding results shown in Fig. 6(a). Figure 6(b)-6(d) are intensity profiles plotted through the center of each bead in three directions. To get more accurate intensity profiles, the intensity data of all the three beads in each direction was fitted to Gaussian curve. The FWHM of each intensity profile could be obtained from the fitted Gaussian curve, which are 6.5 µm, 7 µm and 41 µm in x, y and z directions, respectively.

**Fig. 6.**
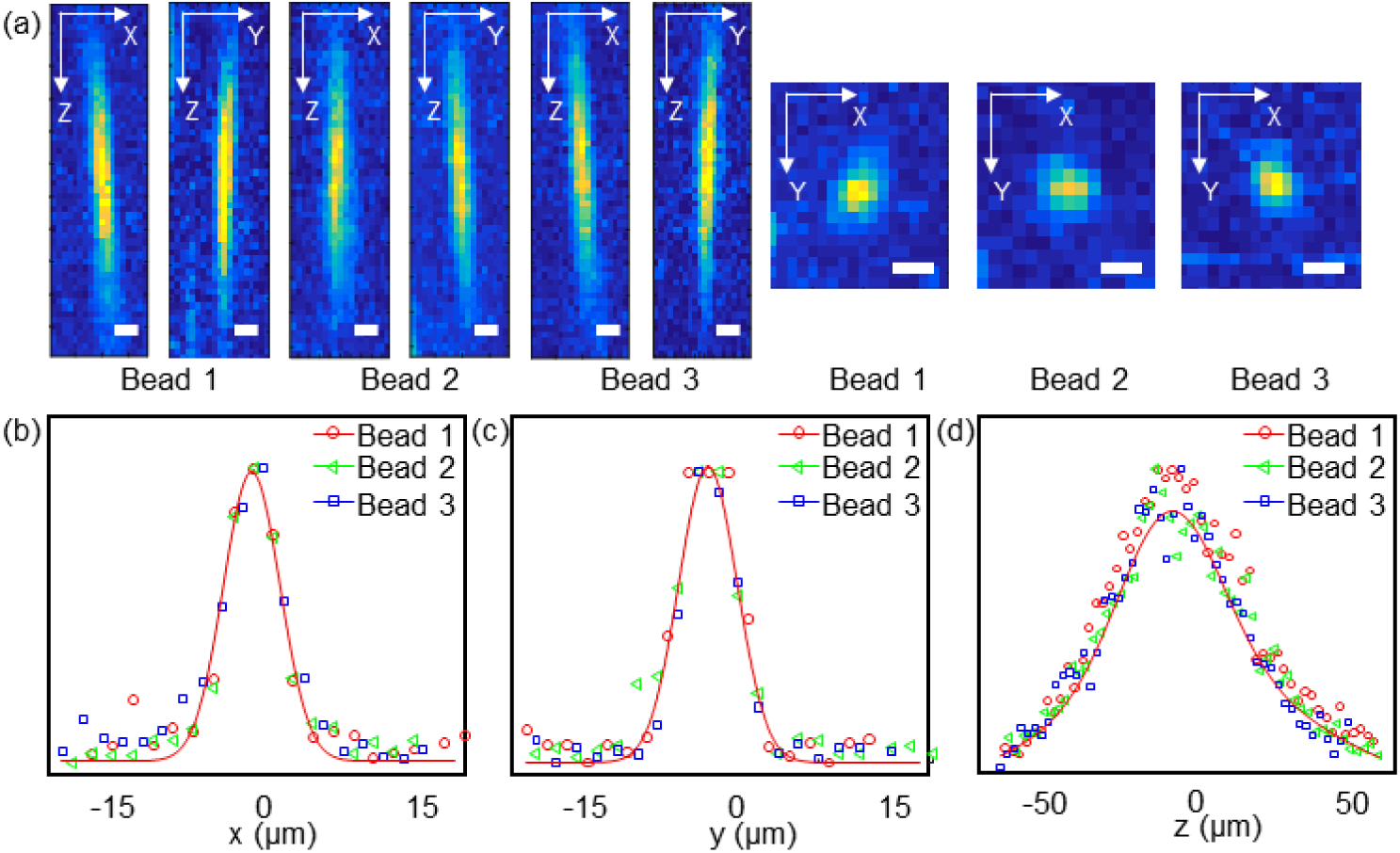
Intensity profile of the three representative beads. (a) Maximum intensity projection of three representative beads along x, y, and z directions with their locations marked in panel (d-f) of Fig. 5. Bar = 10 µm. (a-c) The intensity profile through the center of the three beads in x, y, and z direction.

Fluorescein angiography (FA) is a major retinal imaging modality intended to visualize the vascular leakage by intravenous fluorescein injection. One primary method for FA is SLO that provides 2D images of fluorescein signals. To offer 3D information, we recently achieved volumetric FA (vFA) in rodents’ retina over ∼30° FOV using oSLO, which is to be implemented in human retina with the setup proposed in this paper. In order to evaluate the depth discrimination of oSLO in human retina for vFA, we performed an simulation using *in vivo* OCTA at macular region as the gold standard for the microvasculature, and performed one dimensional convolution to the 3D OCTA image to match the axial resolution of the current oSLO system (41 μm), as shown in Fig. 7(a). We assumed that the convoluted images could be served as a good prediction for vFA in human retina by oSLO. We then applied maximum intensity projection to three segmented layers, as indicated in Fig. 7(b)-7(d). Although the depth resolution of the oSLO is worse than that of OCTA, the three vascular layers in the inner retina could still be separated, as well as the retina circulation and choroidal circulation. Therefore, a similar depth discrimination ability should be achieved by oSLO when it is applied to the living human retina.

**Fig. 7.**
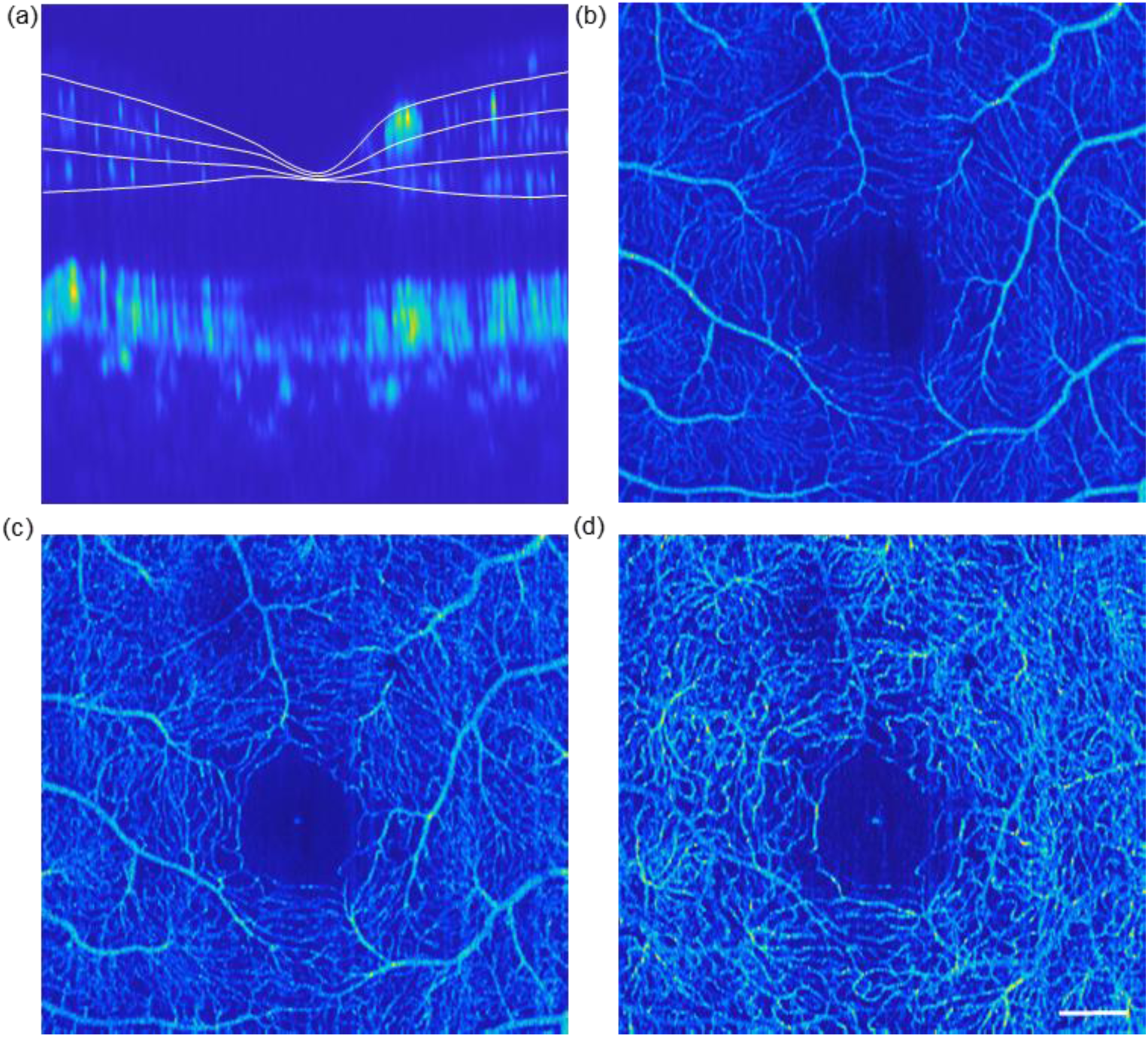
Simulation of the depth discrimination capability of oSLO in human retina based on OCTA. (a) The OCTA cross-sectional image with convoluted depth dimension. (b-d) The *en face* view of retinal in three different layers. Bar = 500 µm.

## 4. Discussion and conclusion

We have proposed a new oSLO system for human retina volumetric fluorescence imaging and have assessed its feasibility and resolution using a realistic eye model. Both the optical property as well as the artificial retina are similar to that of a real human eye. The dimension of the artificial retina is precisely measured by OCT. The resolution of oSLO was assessed to be 6.5 µm, 7 µm and 41 µm in *x, y* and *z* directions, respectively, over a FOV of 2.8 × 6 × 0.8 mm in *x-y-z*. The fast scan *x* dimension is currently limited by the size of the camera sensor and magnification, which can be improved by a larger sensor size and proper magnification. With 10 ms exposure time, the current frame rate could reach 100 fps. By limiting the power of the incident laser beam to 0.5 mW, the imaging procedure is safe for the human eye and compliant to laser safety standard. Therefore, we conclude that the proposed oSLO is feasible for volumetric fluorescence imaging in human retina, providing depth resolution that has been challenging for all the existing fluorescence-based retinal imaging modalities.

The design of the magnification ratio in the detection path, as well as the adoption of cylindrical lenses, overcome the challenge raised by low NA in the human eye in two aspects. Firstly, the tilted image can be increased from 10° to 41° allowing enough space for subsequent optical components; secondly, as shown in Fig. 2, the heavy compression in the axial direction is optically expanded, and the overall symmetrical magnification is achieved. Due to the small NA of the human eye (equivalent to a small NA objective lens in a microscopic setting), the depth resolution of oSLO is fundamentally limited at several tens of microns [28]. Nevertheless, we demonstrated a depth resolution ∼41μm without correcting any aberrations, sufficient to separate retinal circulations by different layers, and from choroidal circulation (Fig. 7).

There are several revenues to further improve our human eye oSLO system. Firstly, the efficiency of the fluorescent collection is limited by the tilted objective lens OL2 in the proposed image system. Applying water immersion objective lens OL2, such as the method described in oblique illumination microscopy [35], would improve the efficiency of the fluorescent collection. Secondly, although 10ms exposure time was demonstrated in the proposed method, the frame rate was limited by the data transfer speed of the currently used camera. This speed can be dramatically improved by adopting a high-speed scientific CMOS camera. Thirdly, our system was built up with off-the-shelf lenses and suffered from the aberration of the human eye such that it did not reach the theoretical limit. Improvements can be achieved by optimizing the off-axis performance and correcting human eye aberration. Finally, all images are shown as acquired, without reconstruction and deconvolution. Additional image processing, such as model-based or inter-plane deconvolution, can be used to improve the resolution [36].

To conclude, a novel oSLO for the human eye, which is capable of volumetric fluorescence retinal imaging, is presented for the first time. The depth resolution is very completive with traditional SLO. The feasibility of the proposed method is demonstrated under a human eye model showing great potential for clinical application.

## References and links

[1]. H. R. Novotny and D. L. Alvis, “A method of photographing fluorescence in circulating blood in the human retina,” Circulation 24, 82–86 (1961).

[2]. P. A. Liebman and R. A. Leigh, “Autofluorescence of Visual Receptors,” Nature 221(5187), 1249–1251 (1969).

[3]. G. K. Frampton, N. Kalita, L. Payne, J. L. Colquitt, E. Loveman, S. M. Downes, and A. J. Lotery, “Fundus autofluorescence imaging: systematic review of test accuracy for the diagnosis and monitoring of retinal conditions,” Eye 31(7), 995–1007 (2017).

[4]. S. J. Ryan, S. R. Sadda, D. R. Hinton, and A. P. Schachat, Retina (2013).

[5]. D. Huang, E. A. Swanson, C. P. Lin, J. S. Schuman, W. G. Stinson, W. Chang, M. R. Hee, T. Flotte, K. Gregory, C. A. Puliafito, and A. Et, “Optical coherence tomography,” Science 254(5035), 1178–1181 (1991).

[6]. Y. Jia, S. T. Bailey, T. S. Hwang, S. M. McClintic, S. S. Gao, M. E. Pennesi, C. J. Flaxel, A. K. Lauer, D. J. Wilson, J. Hornegger, J. G. Fujimoto, and D. Huang, “Quantitative optical coherence tomography angiography of vascular abnormalities in the living human eye,” Proc. Natl. Acad. Sci. 112(18), E2395–E2402 (2015).

[7]. S. Mo, B. Krawitz, E. Efstathiadis, L. Geyman, R. Weitz, T. Y. P. Chui, J. Carroll, A. Dubra, and R. B. Rosen, “Imaging Foveal Microvasculature: Optical Coherence Tomography Angiography Versus Adaptive Optics Scanning Light Ophthalmoscope Fluorescein Angiography,” Invest. Ophthalmol. Vis. Sci. 57(9), OCT130–OCT140 (2016).

[8]. P. Zhang, A. Zam, Y. Jian, X. Wang, Y. Li, K. S. Lam, M. E. Burns, M. V. Sarunic, E. N. P. Jr, and R. J. Zawadzki, “In vivo wide-field multispectral scanning laser ophthalmoscopy–optical coherence tomography mouse retinal imager: longitudinal imaging of ganglion cells, microglia, and Müller glia, and mapping of the mouse retinal and choroidal vasculature,” J. Biomed. Opt. 20(12), 126005 (2015).

[9]. R. H. Webb, G. W. Hughes, and F. C. Delori, “Confocal scanning laser ophthalmoscope,” Appl. Opt. 26(8), 1492–1499 (1987).

[10]. K. V. Vienola, M. Damodaran, B. Braaf, K. A. Vermeer, and J. F. de Boer, “Parallel line scanning ophthalmoscope for retinal imaging,” Opt. Lett. 40(22), 5335–5338 (2015).

[11]. W. J. Donnelly and A. Roorda, “Optimal pupil size in the human eye for axial resolution,” JOSA A 20(11), 2010–2015 (2003).

[12]. J. J. Hunter, C. J. Cookson, M. L. Kisilak, J. M. Bueno, and M. C. W. Campbell, “Characterizing image quality in a scanning laser ophthalmoscope with differing pinholes and induced scattered light,” JOSA A 24(5), 1284–1295 (2007).

[13]. K. Venkateswaran, A. Roorda, and F. Romero-Borja, “Theoretical modeling and evaluation of the axial resolution of the adaptive optics scanning laser ophthalmoscope,” J. Biomed. Opt. 9(1), 132–138 (2004).

[14]. S. A. Burns, A. E. Elsner, K. A. Sapoznik, R. L. Warner, and T. J. Gast, “Adaptive optics imaging of the human retina,” Prog. Retin. Eye Res. 68, 1–30 (2019).

[15]. D. C. Chen, S. M. Jones, D. A. Silva, and S. S. Olivier, “High-resolution adaptive optics scanning laser ophthalmoscope with dual deformable mirrors,” JOSA A 24(5), 1305–1312 (2007).

[16]. T. Y. P. Chui, D. A. VanNasdale, and S. A. Burns, “The use of forward scatter to improve retinal vascular imaging with an adaptive optics scanning laser ophthalmoscope,” Biomed. Opt. Express 3(10), 2537–2549 (2012).

[17]. L. K. Young, T. J. Morris, C. D. Saunter, and H. E. Smithson, “Compact, modular and in-plane AOSLO for high-resolution retinal imaging,” Biomed. Opt. Express 9(9), 4275–4293 (2018).

[18]. P. Mecê, E. Gofas-Salas, K. Grieve, C. Petit, F. Cassaing, J. Sahel, M. Paques, and S. Meimon, “Visualizing and enhancing axial resolution in nonconfocal adaptive optics ophthalmoscopy,” in Ophthalmic Technologies XXIX (International Society for Optics and Photonics, 2019), 10858, p. 108580P.

[19]. R. J. Zawadzki, P. Zhang, A. Zam, E. B. Miller, M. Goswami, X. Wang, R. S. Jonnal, S.-H. Lee, D. Y. Kim, J. G. Flannery, J. S. Werner, M. E. Burns, and E. N. Pugh, “Adaptive-optics SLO imaging combined with widefield OCT and SLO enables precise 3D localization of fluorescent cells in the mouse retina,” Biomed. Opt. Express 6(6), 2191–2210 (2015).

[20]. Y. Geng, A. Dubra, L. Yin, W. H. Merigan, R. Sharma, R. T. Libby, and D. R. Williams, “Adaptive optics retinal imaging in the living mouse eye,” Biomed. Opt. Express 3(4), 715–734 (2012).

[21]. F. Felberer, J.-S. Kroisamer, B. Baumann, S. Zotter, U. Schmidt-Erfurth, C. K. Hitzenberger, and M. Pircher, “Adaptive optics SLO/OCT for 3D imaging of human photoreceptors in vivo,” Biomed. Opt. Express 5(2), 439–456 (2014).

[22]. M. Yamaguchi, S. Nakao, Y. Kaizu, Y. Kobayashi, T. Nakama, M. Arima, S. Yoshida, Y. Oshima, A. Takeda, Y. Ikeda, S. Mukai, T. Ishibashi, and K. Sonoda, “High-Resolution Imaging by Adaptive Optics Scanning Laser Ophthalmoscopy Reveals Two Morphologically Distinct Types of Retinal Hard Exudates,” Sci. Rep. 6(1), 1–14 (2016).

[23]. M. Laslandes, M. Salas, C. K. Hitzenberger, and M. Pircher, “Increasing the field of view of adaptive optics scanning laser ophthalmoscopy,” Biomed. Opt. Express 8(11), 4811–4826 (2017).

[24]. L. Zhang, W. Song, D. Shao, S. Zhang, M. Desai, S. Ness, S. Roy, and J. Yi, “Volumetric fluorescence retinal imaging in vivo over a 30-degree field of view by oblique scanning laser ophthalmoscopy (oSLO),” Biomed. Opt. Express 9(1), 25–40 (2018).

[25]. W. Song, L. Zhou, J. Yi, and J. Yi, “Volumetric fluorescein angiography (vFA) by oblique scanning laser ophthalmoscopy in mouse retina at 200 B-scans per second,” Biomed. Opt. Express 10(9), 4907–4918 (2019).

[26]. American National Standard for Safe Use of Lasers Z136 (The Laser Institute of America, 2007).

[27]. Europe and International Laser/LED Product Safety Regulations: IEC 60825-1:1993+A1:1997+A2:2001 and IEC 60825-1:2007 (2017).

[28]. L. Zhang, A. Capilla, W. Song, G. Mostoslavsky, and J. Yi, “Oblique scanning laser microscopy for simultaneously volumetric structural and molecular imaging using only one raster scan,” Sci. Rep. 7(1), 1–11 (2017).

[29]. J. E. Greivenkamp, Field Guide to Geometrical Optics (SPIE press, 2004).

[30]. W. Song, S. Fu, S. Song, S. Zhang, L. Zhang, S. Ness, M. Desai, and J. Yi, “Longitudinal detection of retinal alterations by visible and near-infrared optical coherence tomography in a dexamethasone-induced ocular hypertension mouse model,” Neurophotonics 6(4), 041103 (2019).

[31]. W. Song, L. Zhou, K. L. Kot, H. Fan, J. Han, and J. Yi, “Measurement of flow-mediated dilation of mouse femoral artery in vivo by optical coherence tomography,” J. Biophotonics 11(11), e201800053 (2018).

[32]. M. Wojtkowski, V. J. Srinivasan, T. H. Ko, J. G. Fujimoto, A. Kowalczyk, and J. S. Duker, “Ultrahigh-resolution, high-speed, Fourier domain optical coherence tomography and methods for dispersion compensation,” Opt. Express 12(11), 2404–2422 (2004).

[33]. Y. Jia, O. Tan, J. Tokayer, B. Potsaid, Y. Wang, J. J. Liu, M. F. Kraus, H. Subhash, J. G. Fujimoto, J. Hornegger, and D. Huang, “Split-spectrum amplitude-decorrelation angiography with optical coherence tomography,” Opt. Express 20(4), 4710–4725 (2012).

[34]. S. Chen, J. Yi, and H. F. Zhang, “Measuring oxygen saturation in retinal and choroidal circulations in rats using visible light optical coherence tomography angiography,” Biomed. Opt. Express 6(8), 2840–2853 (2015).

[35]. B. Yang, X. Chen, Y. Wang, S. Feng, V. Pessino, N. Stuurman, N. H. Cho, K. W. Cheng, S. J. Lord, L. Xu, D. Xie, R. D. Mullins, M. D. Leonetti, and B. Huang, “Epi-illumination SPIM for volumetric imaging with high spatial-temporal resolution,” Nat. Methods 16(6), 501–504 (2019).

[36]. E. M. C. Hillman, D. A. Boas, A. M. Dale, and A. K. Dunn, “Laminar optical tomography: demonstration of millimeter-scale depth-resolved imaging in turbid media,” Opt. Lett. 29(14), 1650–1652 (2004).

